# Macrophages in the reticuloendothelial system inhibit the propagation phase of mouse apolipoprotein A-II amyloidosis

**DOI:** 10.1101/2021.08.18.456782

**Authors:** Hiroki Miyahara, Jian Dai, Ying Li, Cui Xiaoran, Hibiki Takeuchi, Naomi Hachiya, Fuyuki Kametani, Masahide Yazaki, Masayuki Mori, Keiichi Higuchi

## Abstract

Amyloidosis refers to a group of degenerative diseases that are characterized by the deposition of misfolded protein fibrils in various organs. Deposited amyloid may be removed by a phagocyte-dependent innate immune system; however, the precise mechanisms during disease progression remain unclear. We herein investigated the properties of macrophages that contribute to amyloid degradation and disease progression using transmissible apolipoprotein A-II amyloidosis model mice. Intravenously injected AApoAII amyloid was efficiently engulfed by reticuloendothelial macrophages in the liver and spleen and disappeared by 24 h. While cultured murine macrophages degraded AApoAII via the endosomal-lysosomal pathway, AApoAII fibrils reduced cell viability and phagocytic capacity. Furthermore, the depletion of reticuloendothelial macrophages prior to the induction of AApoAII markedly increased hepatic and splenic AApoAII deposition. These results highlight the physiological role of reticuloendothelial macrophages against inter-individual amyloid propagation and suggest the maintenance of phagocytic integrity as a therapeutic strategy to inhibit disease progression.

## Introduction

Amyloidosis is a degenerative disease that is characterized by extracellular deposits composed of pathogenic fibrillar proteins and associated compounds, such as proteoglycans and lipids. Forty amyloidogenic proteins have been identified as the main components of plaques and intracellular inclusions in amyloid-associated disorders, such as Alzheimer’s disease (AD), Parkinson’s disease, senile systemic transthyretin amyloidosis, and systemic AA amyloidosis (Benson *et al*, 2020; Chiti & Dobson, 2017; Iadanza *et al*, 2018). The increased synthesis and reduced clearance of pathogenic amyloid proteins with aging, genetic mutations, inflammation, dialysis, and obesity have a negative impact on protein homeostasis and have frequently been implicated in the development of amyloidosis (Benson *et al*., 2020; Chiti & Dobson, 2017; Gorevic *et al*, 1986; Iadanza *et al*., 2018). The general pathological feature of amyloidosis is progressive degeneration with tissue damage, with the dissociation of deposited amyloid rarely being reported. However, long-term follow-up clinical studies recently showed the regression of amyloid deposits after reductions in circulating precursor proteins by surgical or pharmacological treatments (Gillmore *et al*, 2001; Holmgren *et al*, 1993; Lachmann *et al*, 2007). Spontaneous amyloid regression was observed in experimental AA amyloidosis mice after the discontinuation of inflammatory stimuli (Hawkins & Pepys, 1990; Muhammad *et al*, 2015). These raise a possibility that amyloid deposits may be efficiently cleared by inherent amyloid clearance systems.

Phagocytes, such as macrophages and microglial cells, are specialized cells that respond to pathogens and injury from cell debris, modified lipoproteins, protein deposits, and other harmful substrates. Phagocyte-derived inflammatory and immunological responses frequently occur in close proximity to amyloid deposits, suggesting a central role in the clearance of amyloid deposits (Bodin *et al*, 2010; Nyström & Westermark, 2012). Clinical trials on patients with AD reported that passive immunotherapy using monoclonal anti-amyloid β (Aβ) successfully cleared senile plaques in the brain, and examinations of its clinical benefits on cognitive decline are ongoing (Laversenne *et al*, 2020; Panza *et al*, 2019). In mouse AA amyloidosis, an immunotherapeutic approach using anti-serum amyloid P component (SAP), which is a novel amyloid signature protein, led to the marked regression of hepatic and splenic amyloid deposition, and this effect was eliminated by the pharmacological treatment of macrophage depletion (Bodin *et al*., 2010). In contrast, drug-induced macrophage depletion has been shown to delay the progression of the mouse AA amyloidosis pathology (Kennel *et al*, 2014; Lundmark *et al*, 2013). Activated microglia and bone marrow-derived mononuclear phagocytes are involved in the initiation and progression of AD (Halle *et al*, 2008; Tejera *et al*, 2019). Therefore, it currently remains controversial whether phagocytes contribute to amyloid clearance or disease progression. Moreover, since the pathogenesis of amyloidosis is predominantly attributed to the aggregation and propagation of misfolded proteins, the contribution of phagocytes during the propagation phase is limited due to the lack of a suitable model system.

To address these issues, we investigated phagocyte-related amyloid clearance mechanisms using a transmissible mouse AApoAII amyloidosis model. AApoAII amyloidosis is characterized by systemic amyloid deposition derived from circulating apolipoprotein A-II (ApoA-II), which is a secondary abundant apoprotein of high-density lipoproteins (HDL). In humans, 4 different point mutations, all of which are located in the stop codon of the *APOA2* gene, resulted in variant ApoA-II with a C-terminal 21-residue elongation (Benson *et al*, 2001; Prokaeva *et al*, 2017; Yazaki *et al*, 2001), and one case of wild-type ApoA-II-derived amyloidosis has been reported (Morizane *et al*, 2011). In mice, AApoAII amyloidosis was initially detected in a senescence accelerated-mouse prone model (Higuchi *et al*, 1986). We subsequently reported that laboratory inbred strain C57BL/6 and crossbred strain BDF1 mice spontaneously developed AApoAII amyloidosis with aging (Korenaga *et al*, 2004). There are seven alleles of the *Apoa2* gene among inbred laboratory strains, and the mouse strain with the type C allele of *Apoa2* (*Apoa2^c^*) exhibits accelerated spontaneous AApoAII amyloidosis (Kitagawa *et al*, 2003). We established a congenic strain of mice with the amyloidogenic *Apoa2^c^* allele on the genetic background of the senescence-accelerated resistant mouse (SAMR1), named R1.P1-*Apoa2^c^* (Higuchi *et al*, 1998). The lag phase of AApoAII progression in R1.P1-*Apoa2* mice may be markedly shortened by the administration of a tissue-extracted AApoAII fibril fraction (Higuchi *et al*., 1998). Therefore, R1.P1-*Apoa2^c^* mice represent a valid model for elucidating the cellular and molecular mechanisms of action of phagocytes in both the propagation and progression of amyloidosis.

## Results

### Clearance of injected AApoAII fibrils by hepatic and splenic macrophages *in vivo*

AApoAII fibrils extracted from severely AApoAII-deposited mouse livers using Pras’s method with some modifications (Pras *et al*, 1968) were examined in the present study. This fraction was heterogeneously composed of not only ApoA-II, but also several amyloid signature proteins, such as apolipoprotein E (ApoE), clusterin, and vitronectin (Miyahara *et al*, 2018) (**Fig 1A**). To investigate the contribution of phagocytes to amyloid clearance, we intravenously injected the AApoAII fraction into ApoA-II knockout mice to avoid labeling endogenous circulating ApoA-II, and histologically analyzed the fate of AApoAII (**Fig 1B**). Two hours after the injection, we detected ApoA-II-positive spots in the liver and spleen by immunostaining. In the liver, injected AApoAII associated with non-hepatocyte cells, which were distributed around sinusoids with irregularly shaped nuclei (**Fig 1C**). These cells were positively stained by the macrophage marker F4/80 with ApoA-II signals in the cytoplasm, indicating that injected AApoAII was engulfed by Kupffer cells (**Fig 1D**). We quantitatively confirmed a time-dependent decrease in AApoAII-laden Kupffer cells from 77.0 ± 0.89% to 4.7 ± 1.16% from 2 to 24 h, respectively, after the injection (**Fig 1E and 1F**). Splenic macrophages in the marginal zone also engulfed injected AApoAII (**Fig EV1A**).

**Figure 1.**
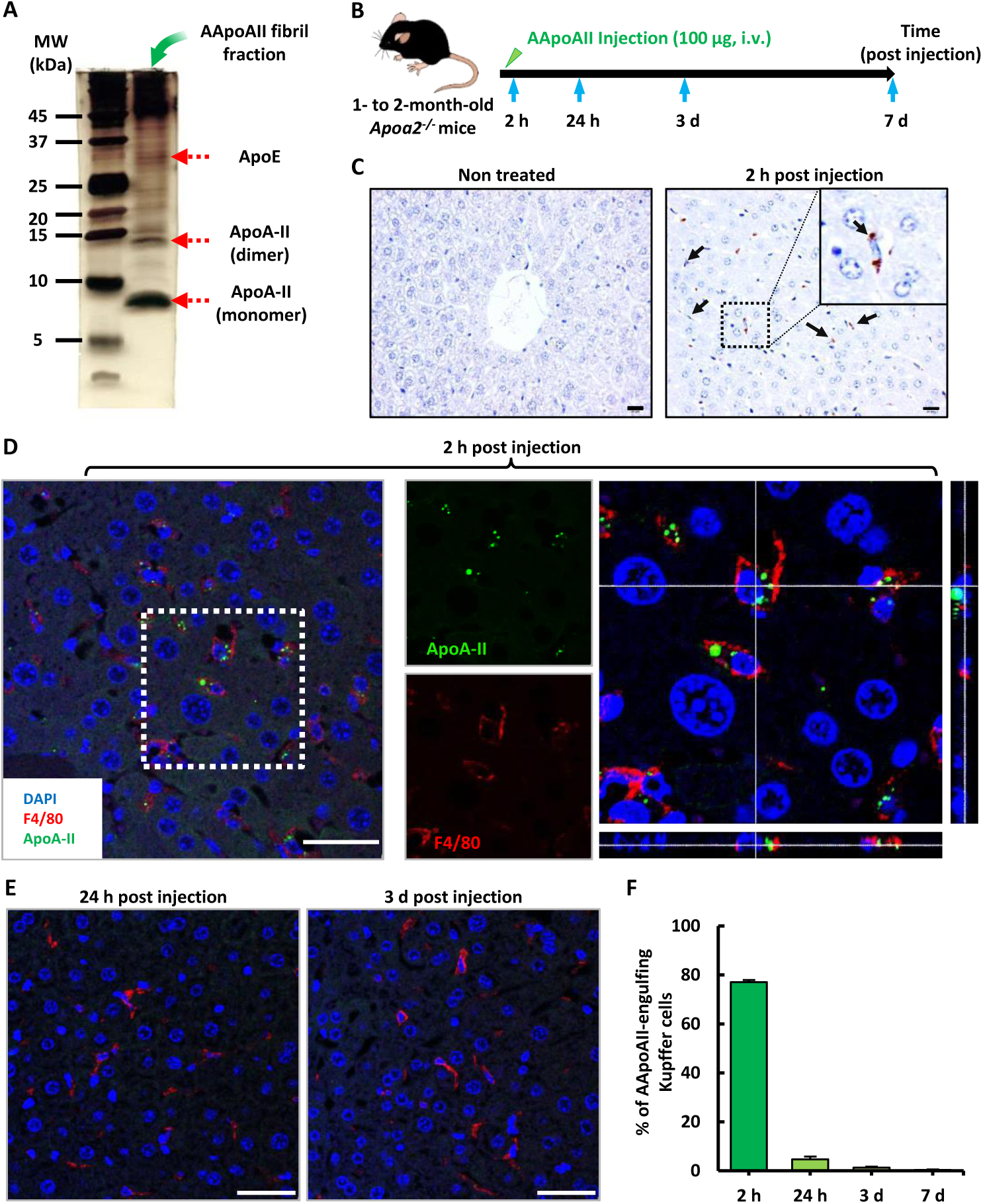
**Engulfment of injected AApoAII fibrils by F4/80-positive Kupffer cells** A: Silver staining of the tissue-extracted AApoAII fibril fraction. The molecular weight marker (left lane) and 2.5 µg of AApoAII (right lane) were separated by SDS-PAGE. B: Experimental design to study the phagocytic contribution to the degradation of AApoAII. B6-*Apoa2^-/-^* mice were euthanized 2 h, 24 h, 3 d, and 7 d after the injection of the AApoAII fraction (n = 5). C: Immunohistochemical staining of liver sections of non-treated control mice (left) and 2 h after the injection (right) using the ApoA-II antibody. Arrows in the boxed region indicate the engulfment of AApoAII by irregularly shaped cells. Scale bars; 20 µm D-F: Immunofluorescence analysis of liver sections of mice (D) 2 h, (E) 24 h, and (F) 3 d after the injection using ApoA-II (green) and F4/80 (red) antibodies. (D) A close-up of the indicated cells (white square) is shown in the right panels. A confocal z-stack image showing F4/80-positive Kupffer cells phagocytosing AApoAII. Scale bars; 20 µm F: Quantitative analysis of the number of AApoAII-engulfing Kupffer cells as indicated in (D-F). Data represent the means ± SEM (standard error of the mean) of samples calculated from 5 fields in each mouse (n = 5).

We then examined the relationship between AApoAII deposits and macrophages using R1.P1-*Apoa2^c^* mice. The R1.P1-*Apoa2^c^* strain is a transmissible mouse model of AApoAII amyloidosis induced by an intravenous injection of AApoAII fibril fractions (Higuchi *et al*., 1998; Miyahara *et al*., 2018). We previously detected hepatic AApoAII deposits in connective tissues and endothelial cells around portal veins 2 months after an injection (Miyahara *et al*., 2018) (**Fig EV1B to E**). F4/80-positive Kupffer cells were distributed in close proximity to AApoAII deposits, possibly reflecting the clearance of AApoAII deposits by hepatic Kupffer cells.

### Cultured macrophages assemble AApoAII fibrils on cell surfaces and degrade under living conditions

To clarify the role of macrophages in the natural clearance system of AApoAII, we investigated whether AApoAII fibrils were degraded in cultured macrophages. The addition of AApoAII fibrils to murine macrophage-like J774A.1 cells led to the formation of cell clusters around AApoAII after 24 h (**Fig 2A**). The presence of AApoAII in the culture medium was monitored using the amyloid fluorescent probe thioflavin T (ThT) (Sawashita *et al*, 2015). In comparisons with immediately after the addition of AApoAII (0 h), the amount of AApoAII in the culture medium decreased with time, with approximately 60% of AApoAII being removed after 24 h (**Fig 2B**). This reduction was not observed in wells without cells or with fixed cells, indicating the amyloid clearance properties of live cells (**Fig 2C**). Transmission electron microscopy (TEM) images confirmed that supplemented AApoAII fibrils adhered to the cell membrane and pseudopodia extended around AApoAII fibrils to engulf them (**Fig EV2**). To investigate the fate of cell-adhered AApoAII fibrils, cells incubated with AApoAII fibrils for 24 h were further incubated in fresh culture media for up to 24 h. We observed a time-dependent decrease in AApoAII signals, the level of which reached 1.4 ± 0.14% by 24 h, and a shift in fluorescence puncta into the cytoplasm (**Fig 2D**).

**Figure 2.**
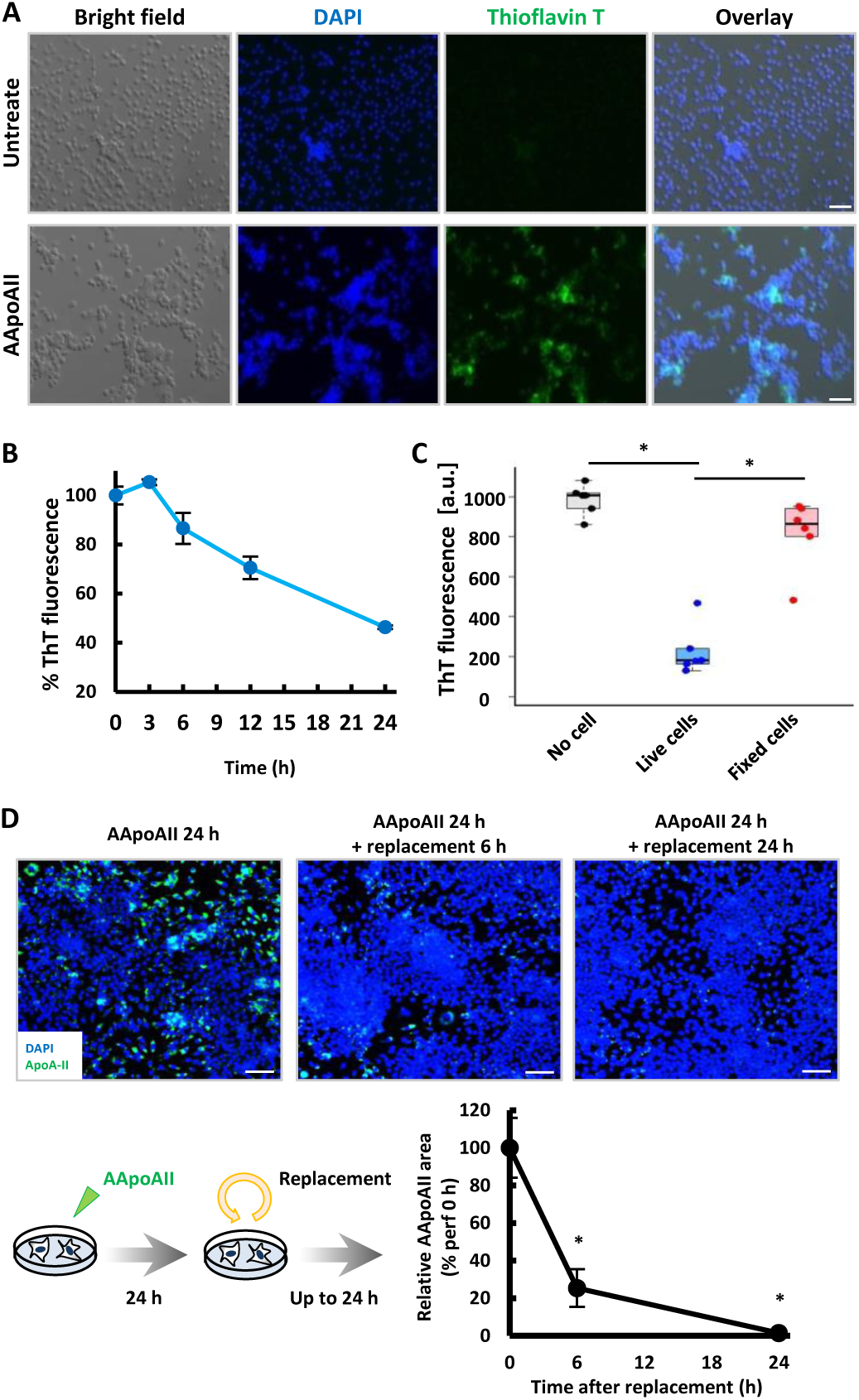
**Tissue-extracted AApoAII amyloid is degraded by cultured macrophages.** A: Representative images of J774A.1 cells incubated with AApoAII at 100 µg/mL for 24 h and stained by thioflavin T (ThT) to visualize AApoAII. Arrows indicate clustered macrophages around AApoAII. Scale bars; 100 µm. B,C: Quantitative analysis of AApoAII remaining in culture media after the co-incubation with AApoAII at 40 µg/mL and B) J774A.1 cells for up to 24 h, C) live cells, fixed cells, or without cells for 24 h. Data represent the means ± SEM of ThT fluorescence in quintuplicate. \**P* <0.05 (Tukey-Kramer method). D: After 24 h of exposure to AApoAII at 40 µg/mL, cells were washed twice and further incubated in fresh culture media for up to 24 h, immunostained with the ApoA-II antibody (green), and cell-associated AApoAII was quantified using ImageJ software. Data represent means ± SEM and five images per well were used for quantification (n = 5). Scale bars; 100 µm. \**P* <0.05 (Tukey-Kramer method).

### Macrophages degrade ApoAII via the endosomal-lysosomal pathway

The results obtained suggested that macrophages degrade extracellular AApoAII by trafficking into intracellular compartments. In some amyloid species, lysosomes are the intracellular organelle responsible for the digestion of toxic aggregates (Kameyama *et al*, 2016; Marshall *et al*, 2020). Therefore, we visualized lysosomes using LysoTracker and examined the co-localization of intracellular AApoAII and lysosomes. After 24 h of AApoAII supplementation, AApoAII fluorescence puncta were observed inside and outside of cells, and co-localized with lysosomes or assembled along extracellular membranes, respectively (**Fig 3A**). When cells were further incubated in culture media without AApoAII, co-localized signals decreased with time. We then examined the impact of endocytic or lysosomal inhibition on intracellular amyloid degradation. The reduction in AApoAII in culture media was significantly attenuated by a pre-treatment with the lysosomal acidic inhibitor bafilomycin A1 (BafA1) and endosomal trafficking inhibitor VPS34-IN1 as well as the actin polymerization-dependent macropinocytosis inhibitor cytochalasin D (**Fig 3B**). In contrast, none of the compounds targeting micropinocytosis, including clathrin-dependent (Pitstop2), caveolin-dependent (Genistein), and dynamin-dependent endocytosis (Dynasore), affected clearance efficiency. We noted that the inhibitors used, except for VPS34-IN1, were not toxic to cells, as confirmed by the MTT assay (**Fig 3C**).

**Figure 3.**
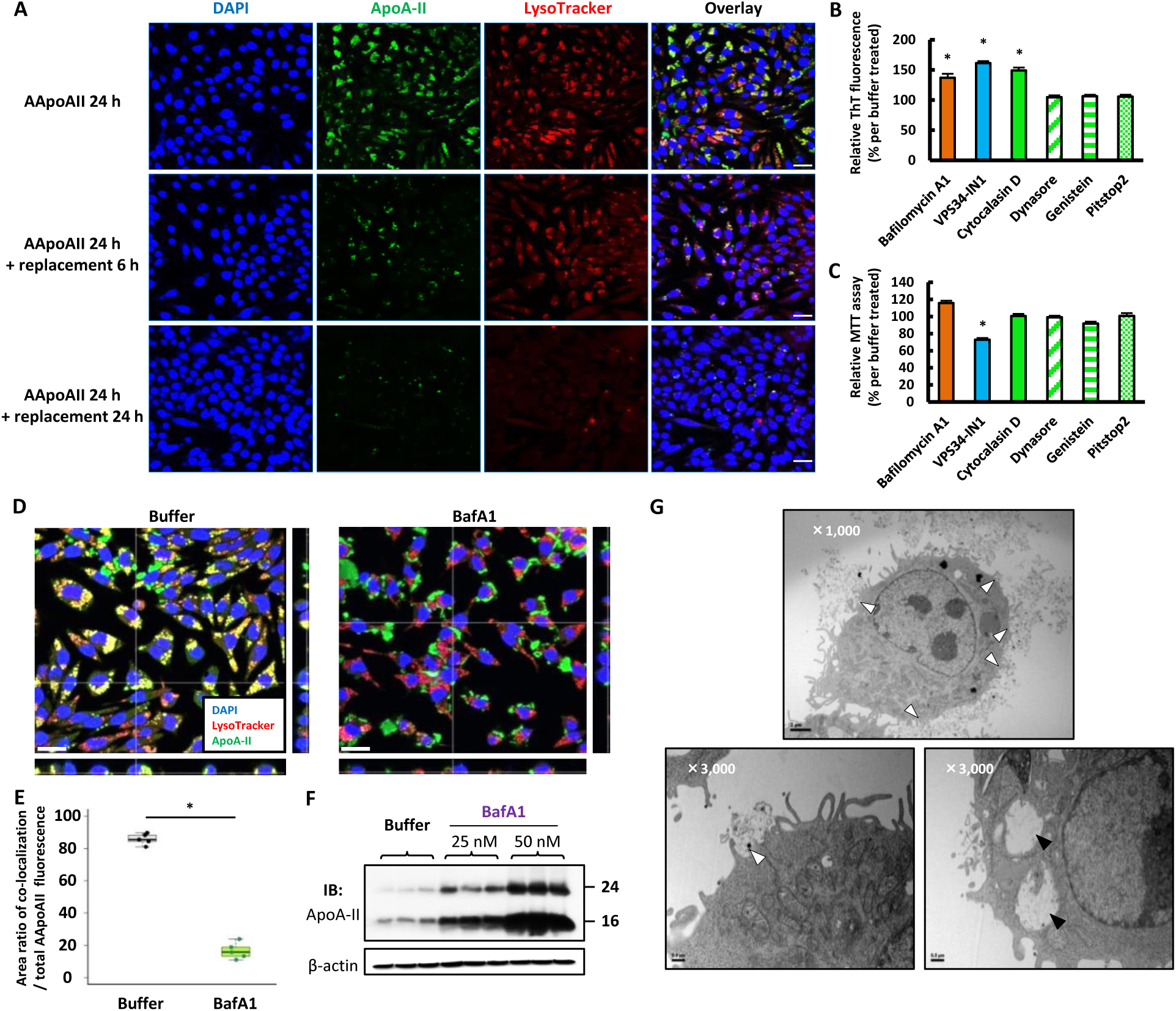
**Cell-associated AApoAII is degraded via the endosomal-lysosomal pathway.** A: Representative confocal images of the co-localization of AApoAII and lysosomes. After a 24-h incubation with AApoAII at 40 µg/mL, J774A.1 cells were further incubated in fresh culture media for up to 24 h, treated with 500 nM LysoTracker (red) for lysosomal labeling for 30 min prior to fixation, followed by immunostaining with ApoA-II (green). Scale bars; 20 µm. B: Quantification of AApoAII remaining in culture media after an incubation with inhibitors using the ThT fluorescence assay. Cells were treated with endocytic and lysosomal inhibitors (Each condition was described in the Methods section), followed by an incubation with AApoAII at 40 µg/mL for 12 h. Values represent the mean ± SEM of the relative percentage to cells without an inhibitor (n = 5). \**P* <0.05 (Tukey-Kramer method). C: MTT-based evaluation of the cytotoxicity of inhibitors. After the treatment with inhibitors, cells were further incubated in culture media for 12 h and used in the MTT assay (mean ± SEM , n = 5). \**P* <0.05 (Tukey-Kramer method). D, E: Immunofluorescence analysis of cells treated with buffer control or bafilomycin A1 (BafA1), followed by an incubation with AApoAII at 40 µg/mL for 24 h, lysosomal labeling (red), and immunostaining by ApoA-II (green). Values represent the mean ± SEM of the relative percentage of the co-localization area (yellow) to total AApoAII fluorescence (n = 5). Scale bars; 20 µm. \**P* <0.05 (the Student’s *t*-test). F: Western blot analysis of cell lysates treated with buffer or BafA1 at 25 or 50 nM for 3 h, followed by an incubation with AApoAII at 40 µg/mL for 24 h. The lower and upper bands of the ApoA-II window indicate dimeric and trimeric ApoA-II, respectively. G: TEM images of cells incubated with AApoAII at 40 µg/mL for 24 h. Higher magnification images indicate the interaction of membrane-adhered AApoAII with extended pseudopods (white arrowheads) and intracellularly endocytosed AApoAII by large endosomes with a diameter > 1 µm (arrowheads). Scale bars; 2 µm (upper) and 0.5 µm (lower).

We demonstrated that macrophages treated with BafA1 showed the delayed degradation of AApoAII amyloid. In comparisons with non-treated cells, the fluorescence signals of co-localization with AApoAII and lysosomes decreased from 85.6 ± 1.50% to 16.7 ± 2.24% (**Fig 3D and E**), with an increase in cell-adhered AApoAII in BafA1-treated cells (**Fig 3F**). This result suggested that lysosomal dysfunction disrupted the uptake of extracellular AApoAII in order to avoid the intracellular accumulation of cytotoxic substrates, which is consistent with previous findings (Xu *et al*, 2003). We observed the co-localization of internalized AApoAII with high-molecular-weight (HMW: 70 kDa) dextran (**Fig EV3**), which is predominantly endocytosed by macropinocytosis (Li *et al*, 2015). TEM images showed the internalization of AApoAII fibrils with membrane ruffles and localization to relatively large endosomes with a diameter of > 1 µm (**Fig 3G**). These results indicated that the engulfment of AApoAII fibrils was mediated by actin polymerization-dependent macropinocytosis, followed by degradation via the endosomal-lysosomal pathway.

### The AApoAII degradation process exhausts the phagocytic capacity

We examined macrophages during the AApoAII degradation process. Previous studies reported that the intracellular accumulation of amyloid fibrils induced cellular damage and pro-inflammatory responses (Friker *et al*, 2020; Yates *et al*, 2000). Based on the MTT assay, significant reductions in cell viability were observed after an incubation with 20 µg/mL AApoAII for 24 h and cytotoxicity increased in a dose-dependent manner (**Fig 4A**). We also investigated whether the intracellular digestion of AApoAII affected the dynamics of macrophages. After 24 h of AApoAII supplementation, we assessed changes in the expression levels of macrophage population markers and apoptosis-related genes using qPCR. AApoAII supplementation induced significant increases in the levels of the alternatively activated macrophage (M2φ) markers *Arg1* and *Il10* over those in buffer control cells (**Fig 4B**). In contrast, the level of the classically activated macrophage (M1φ) marker Il1b decreased. Other macrophage marker genes, such as *Il6*, *Tnf*, and *Tghb1* as well as the apoptosis-related genes, *Bax* and *Bcl2* and lysosome-associated gene, *Lamp2* were not altered by AApoAII supplementation.

**Figure 4.**
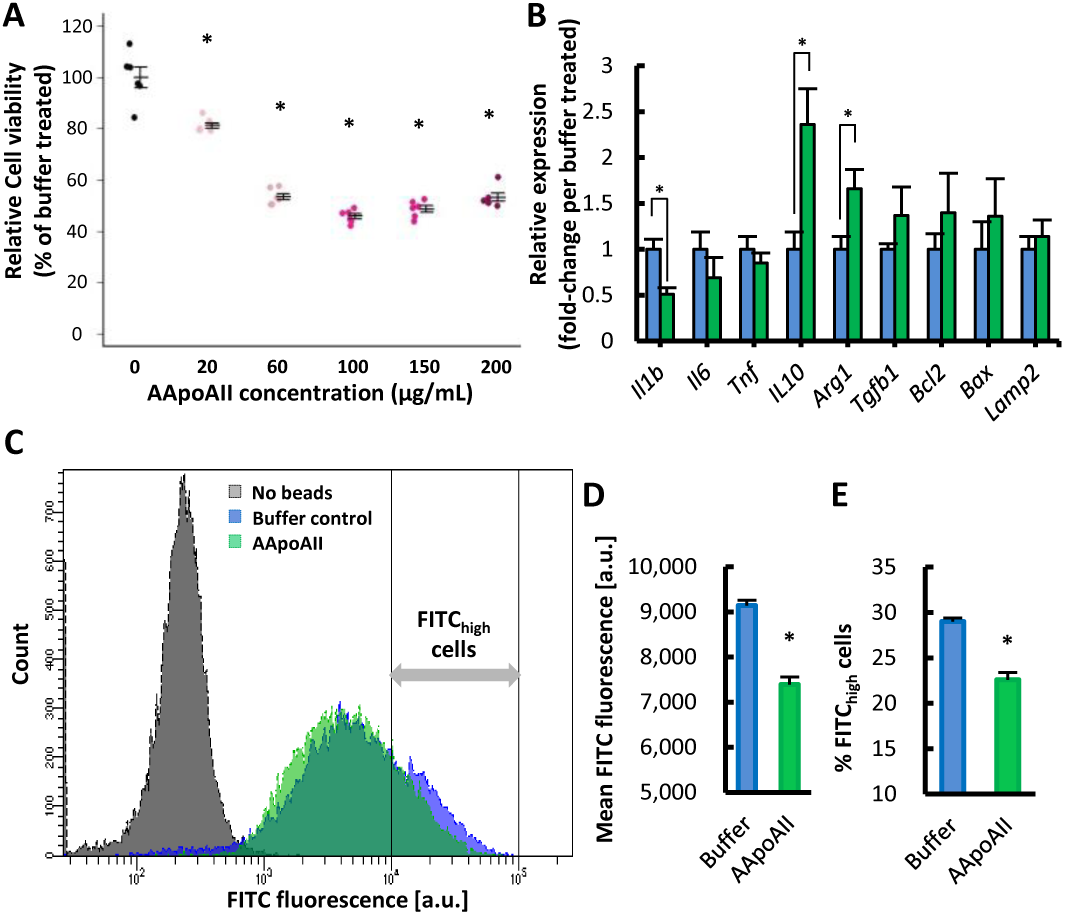
**Exposure to AApoAII induces cytotoxicity and decreased phagocytic capacity.** A: MTT-based cell viability measurements of cells exposed to AApoAII for 24 h (n = 5). Data represent the mean ± SEM (n = 5). \**P* <0.05 (Tukey-Kramer method). B: Quantitative PCR analysis of expression levels of macrophage population markers and apoptosis-related genes in cells incubated with AApoAII at 40 µg/mL for 24 h (n = 5). Data represent the mean ± SEM (n = 5). \**P* <0.05 (the Student’s *t*-test). C-E: Quantification of the phagocytic capacity measured by a flow cytometric analysis of cells incubated with AApoAII at 100 µg/mL for 24 h, followed by an incubation with FITC-labeled beads for 6 h. Data represent the mean ± SEM of (D) FITC fluorescence and (E) the percentage of FITC_High_ cells (n = 3). \**P* <0.05 (the Student’s *t*-test).

We then investigated whether these changes in macrophage dynamics affected the capacity for cellular AApoAII degradation. Since an overloaded lysosomal capacity leads to the exhaustion of proteolytic resources and phagocytic functions in macrophages (Marques *et al*, 2021), we analyzed the phagocytic activity of AApoAII-supplemented cells with FITC-labeled beads using flow cytometry. When cells were supplemented with AApoAII for 24 h followed by the uptake of FITC-labeled beads for 6 h, bead uptake was significantly lower than that by buffer-treated cells (**Fig 4C and D**). Additionally, we found that some buffer-treated macrophages showed a high capacity for bead uptake (FITC_High_ cells) and this population was reduced by the AApoAII treatment (**Fig 4C and E**), suggesting that the surplus phagocytic capacity was exhausted by the AApoAII degradation process.

### Depletion of reticuloendothelial macrophages exacerbate hepatic and splenic AApoAII deposition

Based on the present results showing that macrophages have the potential to degrade AApoAII, we examined the effects of macrophage depletion in the AApoAII amyloidosis pathology. Liposomes containing the cytotoxic drug clodronate (CLO) were selectively phagocytosed by reticuloendothelial cells, which lead to the temporary depletion of hepatic and splenic macrophages (Van Rooijen & Sanders, 1994). R1.P1-*Apoa2^c^* mice were intravenously administered control or CLO liposomes with a diameter of 280 ± 50 nm at 25 mg/kg body weight. After 24 h, F4/80-positive Kupffer cell numbers in the liver were lower in mice treated with CLO liposomes (pre-CLO) than in the control liposome group (pre-control); therefore, these mice were further injected with AApoAII fibrils to induce AApoAII amyloidosis (**Fig 5A**). After 2 months, AApoAII deposits detected by ThT fluorescence in the liver and spleen were markedly higher (10.6- and 2.5-fold, respectively) in pre-CLO mice than in pre-control mice (**Fig 5B and C**). We also investigated the effects of macrophage depletion after the onset of AApoAII amyloidosis. Mice were administered 25 mg/kg body weight CLO or control liposomes one month after induction (post-CLO and post-control), and were euthanized two months after the AApoAII injection (**Fi. 5D and E**). In contrast to the pre-treatment experiment, no changes in AApoAII levels were detected in the liver or spleen between the two groups (**Fig 5F**).

**Figure 5.**
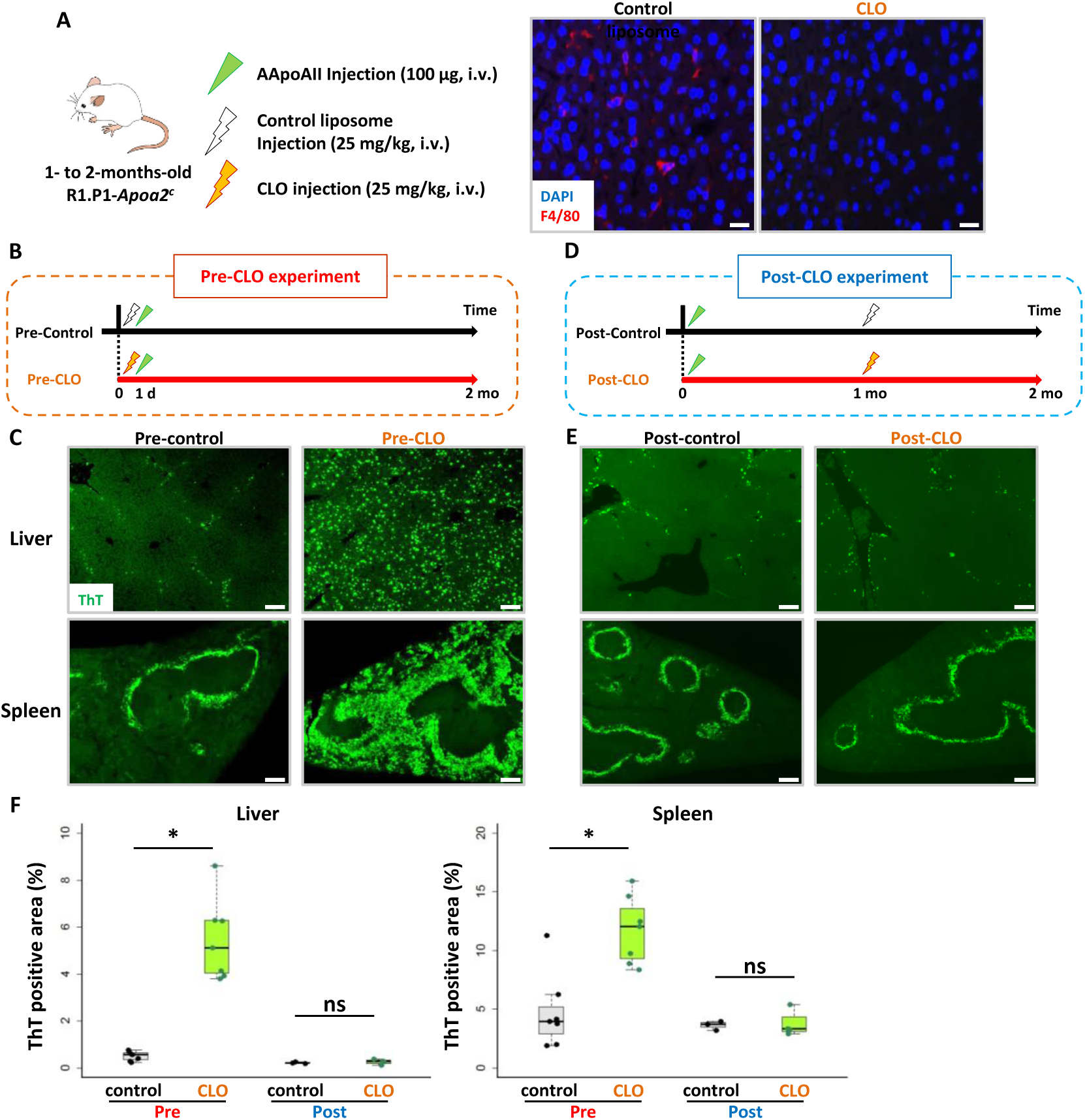
**Depletion of reticuloendothelial macrophages accelerates hepatic and splenic AApoAII deposition.** A: Representative images of hepatic macrophage depletion in R1.P1-*Apoa2^c^* mice treated with CLO at 25 mg/kg for 24 h. Kupffer cells were immunostained by F4/80 (red). Scale bars; 20 µm. B-E: Experimental design of the evaluation of macrophage depletion in the AApoAII amyloidosis pathology. (B) Twenty-four hours after the CLO treatment, mice were injected with 100 µg AApoAII to induce AApoAII amyloidosis and then sacrificed at 2 months (n = 7). (D) One month after the injection of 100 µg AApoAII, mice were treated with CLO and sacrificed at 2 months (n = 3). (C and E) Representative images of hepatic and splenic AApoAII deposition stained by ThT. Scale bars; 200 µm. F: Quantitative analysis of hepatic and splenic ThT-stained AApoAII deposition areas (green) by ImageJ software. Data represent the mean ± SEM. Three images at a magnification of ×5 objective lens per mouse were used for quantification. \**P* <0.05 (the Student’s *t*-test). ns; not significant.

We noted that diffuse amyloid deposition in the liver is a characteristic signature of inflamed systemic mouse AA amyloidosis (Muhammad *et al*., 2015; Westermark & Westermark, 2009). However, serum amyloid A (SAA) was not detected in the amyloid deposits of pre-CLO mice (**Fig EV4A**). In addition, serum levels of ApoA-II remained unchanged from 1 day to 2 months after the CLO treatment (**Fig EV4B**), indicating that the development of AApoAII deposits was independent of AA amyloid and the production of ApoA-II. In contrast to increases in AApoAII deposition in the liver and spleen, other organs, including the lung, heart, stomach, small intestine, skin, and tongue, did not show any pathological changes between pre-CLO and pre-control mice (**Fig EV4C**). Collectively, the present results revealed the AApoAII degradation mechanisms of macrophages, which prevented the propagation of external AApoAII at the reticuloendothelial system.

## Discussion

Numerous clinical studies have revealed that deposited amyloid naturally regresses after pathogenic amyloid protein production has been blocked (Gillmore *et al*., 2001; Holmgren *et al*., 1993; Lachmann *et al*., 2007). Innate immune phagocytic cells were suggested to be the key mediators of the natural amyloid clearance system (Bodin *et al*., 2010). Failure to clear amyloid is assumed to contribute to the pathogenesis of amyloidosis; however, the underlying mechanisms have not yet been elucidated in detail. We herein investigated the natural amyloid clearance mechanisms of macrophages using a transmissible AApoAII amyloidosis model, both *in vivo* and *in vitro*. The results obtained showed that macrophages efficiently degraded tissue-extracted AApoAII amyloid and reticuloendothelial macrophages prevented disease progression, at least in the propagation phase.

We demonstrated the degradation of AApoAII via the endosomal-lysosomal pathway in a murine macrophage cell line (**Fig 3**). Macrophages initially bound AApoAII fibrils on their plasma membrane, followed by their uptake by macropinocytosis and the trafficking of extracellular fibrils into lysosomes. These events mostly occurred within 24 h both *in vivo* and *in vitro*. Similar to the present results, previous studies reported the cellular properties of amyloid degradation. The engulfment of amyloid fibrils and protein aggregates was shown to be mediated by macropinocytosis via heparan sulfate proteoglycans on the cell surface (Holmes *et al*, 2013; Kuwabara *et al*, 2015). The cellular uptake and degradation of amyloid fibrils are reported not only for phagocytes but also parenchymal cells of amyloid-laden tissues (Kameyama *et al*., 2016; Okoshi *et al*, 2015). Importantly, internalization and degradation by lysosomes appear to key events in amyloid clearance, with the intracellular accumulation of amyloid fibrils and overloading of the lysosomal proteolytic capacity triggering the disruption of the endosomal-lysosomal pathway, mitochondrial depolarization, and, ultimately, cell death (Kameyama *et al*., 2016; Marshall *et al*., 2020; Okoshi *et al*., 2015). Accordingly, the present results showed that cultured macrophages reduced their phagocytic capacity during AApoAII degradation (**Fig 4**). Therefore, amyloid fibrils may lead to the negative feedback of these functions in cells with disease progression.

Recent genome-wide association studies on late-onset AD identified genes implicated in the endosomal-lysosomal pathway, such as bridging integrator 1, phosphatidylinositol-binding clathrin assembly protein, and sortilin-related receptor 1 (Lambert *et al*, 2013). The endosomal-lysosomal pathway of phagocytes was found to be impaired during aging and in patients with AD (Solé-Domènech *et al*, 2016; Van Acker *et al*, 2019). Lysosomal dysfunction induced the formation of intracellular amyloid (Claus *et al*, 2017) and the release of exosomes containing APP fragments or toxic Aβ oligomers, which facilitate the cell-to-cell propagation of amyloid (Miranda *et al*, 2018; Sardar Sinha *et al*, 2018). Based on these findings, we propose the maintenance of endosomal-lysosomal integrity as a therapeutic strategy to inhibit disease progression and promote the clearance of amyloid deposits.

The pro-inflammatory factors, granulocyte-macrophage colony-stimulating factors (GM-CSF and M-CSF) have been shown to enhance microglial proliferation and lysosomal activity, and subsequently enhance amyloid clearance properties (Daria *et al*, 2017). Similar beneficial effects have been reported in cerebral amyloid angiopathy model mice stimulated with perivascular macrophages by the administration of chitin (Hawkes & McLaurin, 2009). Moreover, soluble factors released from young microglia restore the amyloid clearance capacity of old microglia even at advanced stages of amyloid deposition (Daria *et al*., 2017). In the present study, the gene expression levels of alternatively activated M2φ markers were higher than those of classically activated M1φ markers during the degradation of AApoAII in cultured macrophages. Based on reductions in cellular viability and the phagocytic capacity, this population appeared to be impaired with amyloid clearance systems. Although our experiments did not allow us to elucidate the mechanisms underlying this outcome, further studies on both phagocytic populations and the factors facilitating amyloid clearance will facilitate the development of beneficial strategies for amyloidosis. However, since the phagocytes activated by pro-inflammatory pathways are directly linked to disease progression (Lucin & Wyss-Coray, 2009), the findings should be interpreted with caution.

The depletion of macrophages in the reticuloendothelial system markedly increased the hepatic and splenic amyloid loads, but not those in other organs, which were not targeted by CLO included by large-scale liposomes (**Fig 5**). Similar to the present results, the depletion of perivascular macrophages was previously shown to increase vascular levels of Aβ (Hawkes & McLaurin, 2009). A study on an *ex vivo* brain model also demonstrated that CLO prevented the clearance of diffuse senile plaques (Daria *et al*., 2017). These findings indicate that phagocytes are a part of the inherent amyloid degradation system; however, macrophage depletion in AA amyloidosis model mice also delayed disease progression (Kennel *et al*., 2014; Lundmark *et al*., 2013). This opposite contribution of macrophages is partially explained by the lysosomal recruitment and processing of soluble proteins by macrophages being involved in the amyloid formation mechanisms of SAA and light chains (Claus *et al*., 2017; Teng *et al*, 2014). Therefore, further studies to clarify the molecular mechanisms underlying amyloid clearance are needed in parallel with those on intracellular amyloid formation.

Unfortunately, the progression of hepatic and splenic AApoAII deposition was not altered in post-CLO mice, indicating the low potential of reticuloendothelial macrophages to clear already deposited AApoAII. Since our *in vitro* results demonstrated that macrophages may degrade AApoAII fibrils, attempts to promote phagocytic functions toward AApoAII deposits may inhibit disease progression. Passive immunization against “amyloid deposits” has been proposed as a putative therapeutic strategy (Bodin *et al*., 2010). Amyloid deposits are generally composed of not only fibrillar proteins, but also some amyloid signature proteins, such as SAP, ApoE, ApoA-IV, clusterin, and vitronectin. Importantly, these amyloid signature proteins are commonly found in various types of amyloidosis regardless of differences between organs and species (Benson *et al*., 2020). Further studies that target amyloid signature proteins may lead to therapeutic advances that are applicable to the different types of amyloidosis, and reductions in circulating precursor amyloidogenic proteins are needed.

Intriguingly, while the hepatic AApoAII pathology of R1.P1-*Apoa2^c^* mice is generally attributed to deposition around portal regions (Miyahara *et al*., 2018), pre-CLO mice showed a different pathology with diffuse amyloid deposition around sinusoids, similar to AA amyloidosis (Westermark & Westermark, 2009). This finding indicates that macrophage depletion induced the progression of AApoAII amyloidosis by alternative mechanisms. Moreover, the distribution of hepatic AApoAII deposition in pre-CLO mice was similar to the biodistribution of intravenously injected AA amyloid trapped by the reticuloendothelial system (Kennel *et al*., 2014). Therefore, we suggest that an alternative pathology resulted from the nucleation-dependent polymerization of trapped amyloid (Naiki *et al*, 2016), which may ordinarily be removed by healthy macrophages. We conclude that reticuloendothelial macrophages are a major protector against individual-to-individual propagation through the efficient clearance of circulating amyloid.

## Materials and Methods

### Animals

R1.P1-*Apoa2^c^* mice are a congenic strain with the amyloidogenic *Apoa2^c^* allele of the senescence-accelerated mouse prone 1 strain on the genetic background of the SAMR1 strain (Higuchi *et al*., 1998). Homozygous ApoA-II knockout (129S4/SvJae-*Apoa2^tm1Bres^*) mice were purchased from Jackson Laboratory and crossbred for 10 generations to a pure C57BL/6 background in our laboratory (B6-*Apoa2^-/-^*). Mice were reared in the Division of Animal Research, Research Center for Support of Advanced Sciences, Shinshu University, under specific pathogen-free conditions at 24 ± 2 °C with a controlled light regimen (12-h light/dark cycle). A commercial diet (MF; Oriental Yeast, Tokyo, Japan) and tap water were available *ad libitum*. All experiments were performed with the approval of the Committee for Animal Experiments at Shinshu University. Between three and seven biological replicates of each mouse sample were used in experiments.

### Preparation of the AApoAII fibril stock

The AApoAII fibril fraction was isolated from the livers of R1.P1-*Apoa2^c^* mice by Pras’s method (Pras *et al*., 1968) with some modifications (Miyahara *et al*., 2018). In brief, 500 mg of a frozen AApoAII-laden liver was homogenized with a Polytron homogenizer (IKA, Baden-Württemberg, Germany) in 5 mL of 0.15 M NaCl, the mixture was centrifuged at 40,000 × *g* at 4°C for 30 min, and the precipitate was collected. This was repeated 4 times. The last precipitate was resuspended in 300 was homogenized in 5 mL of distilled deionized water (DW), the mixture was centrifuged at 30,000 × *g* at 4°C for 30 min, and the supernatant was collected. This was repeated 3 times. The collected solution was ultracentrifuged at 100,000 × *g* at 4°C for 60 min with an ultracentrifuge (Hitachi Koki, Tokyo, Japan). The supernatant was discarded, and the precipitate was resuspended in DW and sonicated with a homogenizer (TIETECH, Nagoya, Japan) on ice for 30 sec. This solution was referred to as the AApoAII fibril fraction. Protein concentrations were estimated using a BCA protein quantification kit (Thermo Fisher Scientific, Tokyo, Japan) according to the manufacturer’s instructions. AApoAII fibrils were stored in a deep freezer at - 80°C until used for experiments.

### AApoAII fibril injections

As we previously reported, the R1.P1-*Apoa2^c^* strain is a transmissible mouse model of AApoAII amyloidosis induced by an injection of AApoAII fibrils (Higuchi *et al*., 1998). One- to two-month-old R1.P1-*Apoa2^c^* or B6-*Apoa2^-/-^* mice were intravenously injected with a single dose of 100 µg of AApoAII fibrils. At each experimental time point, mice were euthanized by cardiac puncture under deep sevoflurane anesthesia. A portion of the tissues was fixed in 10% (v/v) neutral buffered formalin followed by embedding in paraffin for histochemical examinations.

### Histology and immunohistochemistry

Four-micrometer-thick sections of paraffin-embedded tissues were used in the histochemical analysis. Sections were deparaffinized in Hemo-De (FALMA, Tokyo, Japan), rehydrated with ethanol, washed in PBS three times, and blocked by 5% bovine serum albumin (BSA) in PBS at RT for 30 min. Samples were incubated overnight at 4°C in solution containing 1% BSA and primary antibodies: mouse F4/80 (T45-2342, BD Biosciences, New Jersey, USA) 1:200, rabbit antiserum against ApoA-II and SAA prepared in our laboratory 1:1000. Regarding optical microscopic observations, samples were developed using the HRP/DAB staining method, and captured using a Olympus BX43 light microscope and cellSens software (Olympus, Tokyo, Japan). In fluorescence microscopic observations, sections were incubated at RT for 1 h in solution containing appropriate secondary antibodies conjugated to fluorescents: Alexa Fluor 488 (A11008, Invitrogen, Massachusetts, USA) and 555 (A21434, Invitrogen) 1:1000 and NorthernLights NL493 (NL015, R&D Systems, Minnesota, USA) 1:1000, and stained with 4ʹ ,6-diamidino-2-phenylindole (DAPI) for 10 min prior to mounting in Fluoromount^TM^ (K024, Diagnostic BioSystems, California, USA). Samples for ThT staining were treated with 1% ThT for 3 min and differentiated in 1% acetic acid for 15 min. Sections for the double labeling of ThT and ApoA-II/SAA were initially stained using ThT procedures, followed by immunostaining procedures. Sections were washed twice in PBS between each step.

To quantify AApoAII-engulfing Kupffer cells, microphotographs and Z-stack images were acquired using a Leica TCS SP8 confocal microscope with LAS X software. The total numbers of F4/80-positive and AApoAII-engulfing Kupffer cells were manually counted from acquired images and their percentages were calculated. Five images were randomly captured using a ×40 oil objective lens and from each section. To quantify the AApoAII-positive area, microphotographs were acquired using a Zeiss Axio Observer Z1 fluorescence microscope with the AxioVision software package. The fluorescence area of each image was analyzed using ImageJ software (NIH). Three images are randomly captured using a ×5 objective lens from each section.

### Cell culture studies

The murine macrophage-like cell line J774A.1 was purchased from the JCRB Cell Bank. Cells were cultured at 37°C in an atmosphere of 5% CO_2_ using culture media containing RPMI-1640 (FUJIFILM Wako, Osaka, Japan), 10% fetal bovine serum, and 1% penicillin/streptomycin solution. Cell culture medium was replaced every 2-3 days. Cells were plated out at a density of 100,000 cells/mL in 96- or 24-well plates. After 2 days, cells were approximately 80% confluent and used in each experiment. In microscopic analyses, cells were cultured on poly-lysine-coated glass coverslips (Matsunami Glass, Osaka, Japan).

### Immunocytochemistry and labeling of intracellular organelles

Cells were fixed overnight in 4% paraformaldehyde at 4°C and permeabilized in PBS containing 0.5% Triton X-100 at RT for 20 min. After washing with PBS, cells were incubated in blocking solution containing 5% BSA in PBS. Primary and secondary antibodies were sequentially incubated for 1 h in the same buffer, followed by staining with DAPI for 10 min. The coverslip was mounted on a slide grass with Fluoromount^TM^ and imaged using a Leica confocal SP8 confocal microscope or Zeiss Axio Observer Z1 fluorescence microscope.

To visualize lysosomes, cells were treated with fresh medium containing 500 nM LysoTracker Red DND-99 (Invitrogen) for 30 min under growth conditions prior to fixation. In experiments using HMW dextran, cells were incubated in culture media containing both 40 µg/mL AApoAII and 100 µg/mL 70,000-kDa dextran for 24 h prior to fixation.

### ThT fluorescence assay

The amount of AApoAII remaining in culture medium was quantified using a fluorometric assay with ThT. An incubation of 80% confluent cells with medium containing AApoAII at 40 µg/mL was performed. The culture medium was collected and centrifuged at × *g* for 20 min. Pellets were resuspended in 50 µL DW and 20-µL aliquots of each sample were mixed with 180 µL of reaction buffer containing 250 nM ThT and 50 mM glycine-NaOH (pH 9.0). After briefly mixing the solutions at room temperature, ThT fluorescence was analyzed with the Molecular Devices fluorescence spectrometer M5 at excitation and emission wavelengths of 450 and 482 nm, respectively.

### Pharmacological blocking of AApoAII uptake

To investigate the mechanisms underlying AApoAII clearance pathways, cells were incubated with fresh medium containing 50 µM bafilomycin A1 (Adipogen, California, USA), 10 nM VPS34-IN1 (Cayman Chemical), or 2 µM cytochalasin D (Cayman Chemical) for 3 h, or 40 µM Dynasore (Abcam, Cambridgeshire, UK), 20 µM Genistein (Abcam), or 25 µM Pitstop2 (Abcam) for 1 h. After washing, cells were further incubated with fresh medium containing 40 µg/mL of AApoAII for 12 h, followed by the measurement of AApoAII remaining in culture media using the ThT fluorescence assay. All inhibitors were initially dissolved in DMSO as a 1000-fold concentration and stored at -20 °C.

### 3-(4,5-dimethylthiazol-2-yl)-2,5-diphenyltetrazolium bromide (MTT) cell viability assay

To evaluate the cytotoxicity of AApoAII, cells were incubated with fresh medium containing different concentrations of AApoAII for 24 h. After washing, cell viability was assessed using Cell Proliferation Kit I (Roche) according to the manufacturer’s instructions. Absorbance was measured with the Molecular Devices spectrometer M5 at a wavelength of 580 nm.

### Phagocytic capacity assay with a flow cytometer

Phagocytic capacity was assessed using a Phagocytosis Assay Kit (Cayman Chemical, Michigan, USA) according to the manufacturer’s instructions. Cells were treated with AApoAII at 100 µg/mL for 24 h, followed by an incubation in culture media containing FITC-labeled beads for 6 h. Cells were then washed twice with PBS, gently scraped off from each well, and transferred into a 1.5-mL tube. The suspension was centrifuged at 800 × *g* at 4°C for 3 min and the supernatant was discarded. The cell pellet was resuspended into 500 µL of assay buffer and used for a cytometric analysis. Cellular fluorescence was analyzed with a FACSCanto II flow cytometer (BD Bioscience-JP), and data were analyzed using BD FACSDiva software. In each sample, 20,000 events were measured.

### TEM

In TEM experiments, cells grown on coverslips were prefixed in 2.5% (w/v) glutaraldehyde in PBS (pH 7.4) and postfixed in 1% osmium tetroxide, followed by embedding in an epoxy resin block. Ultrathin sections with a thickness of 100-nm were stained with 1% (w/v) uranyl acetate for 10 min followed by an incubation with lead citrate at room temperature for 5 min. Ultramicrographs were acquired using a JEM-1400 electron microscope (JEOL, Tokyo, Japan). The fibrillar structure of AApoAII was detected at ×10,000 magnification.

### Western blot analysis

After washing cells with PBS, the total cell lysate was extracted from cultured cells using RIPA Buffer reagents (Santa Cruz Biotechnology) according to the manufacturer’s instructions. Protein concentrations were assayed using the Pierce BCA Protein Assay Kit (Thermo Scientific, Tokyo, Japan).

In the Western blot analysis, 5-10 µg of total cell lysates was separated by electrophoresis on e-PAGEL Tris-HCl 15% SDS polyacrylamide gels (ATTO, Tokyo, Japan), followed by transfer to polyvinylidene difluoride membranes using the Semi Dry Blotter (ATTO, Tokyo, Japan). Membranes were blocked with 5% non-fat dry milk in Tris-buffered saline at room temperature for 1 h and probed with primary antibodies. Protein bands were detected with the enhanced chemiluminescence method (FUJIFILM Wako) and chemiluminescence film (GE Healthcare, Buckinghamshire, UK), and target protein levels were analyzed using NIH ImageJ software.

### Quantitative PCR analysis

After washing cells with PBS, we extracted total RNA from cultured cells using TRIzol RNA isolation reagents (Invitrogen). Genomic DNA was removed using the DNA-free Kit (Invitrogen). Ten micrograms of total RNA was reverse-transcribed to cDNA using the High Capacity cDNA Reverse Transcriptional Kit with random primers (Invitrogen). Each step was performed according to the manufacturer’s instructions.

A quantitative PCR assay was performed using an ABI PRISM 7500 Sequence Detection System (Applied Biosystems, Tokyo, Japan) with SYBR Green Premix EX Taq^TM^ II (Takara Bio, Tokyo, Japan). *Actb* was used as the internal reference gene. Primers (**Table EV1**) were purchased from Merck Millipore. Each reaction mixture was assayed in duplicate and the average was calculated.

### Depletion of reticuloendothelial macrophages

CLO-containing liposomes with a diameter of 280 ± 50 nm, which selectively target reticuloendothelial macrophages, were purchased from Hygieia Bioscience (http://hygieiabio.co.jp/, Osaka, Japan). One- to two-month-old R1.P1-*Apoa2^c^* mice were intravenously injected with CLO liposomes at 25 mg/kg body weight. The depletion of macrophages was immunohistochemically assessed using the macrophage marker F4/80.

### Statistical analysis

The error bars of each graph represent the standard error (SE). To test the significance of differences, the unpaired Student’s *t*-test for pairwise comparisons and the Tukey-Kramer method for multiple comparisons were performed. *P* values <0.05 were considered to be significant.

## Acknowledgments

The authors thank Dr. Kiyoshi Matsumoto and Dr. Takahiro Yoshizawa (Research Center for Support of Advanced Science, Shinshu University) for animal care. This work was supported in part by the Ministry of Education, Culture, Sports, Science and Technology, Japan (Grant-in-Aid for Scientific Research (B) 17H04063, Research Activity Start-up 19K21258, and Early Career Scientists 20K16215) and the Japan Society for Promotion of Science (Core-to-Core Program A (Advance Research Networks)).

## Author Contributions

H.M., N.H., M.Y., M.M. and K.H. designed research; H.M., J.D., Y.L., C.X. and F.K. performed research; H.M., H.T. and F.K. analyzed data; and H.M. and K.H. wrote the paper.

## Conflict of interest statement

The authors declare that there are no conflicts of interest.

Figure EV1. Relationship between AApoAII and reticuloendothelial macrophages *in vivo*.

A: Immunofluorescence analysis of liver sections of mice 2 h after the injection using ApoA-II (green) and F4/80 (red) antibodies. AApoAII fluorescence was detected in splenic macrophages in the marginal zone (white arrow).

B: R1.P1-*Apoa2^c^* mice were injected with 100 µg AApoAII to induce AApoAII amyloidosis and then sacrificed at 2 months.

C: Immunohistochemical staining of liver sections of non-treated control mice and induced mice using the ApoA-II antibody. AApoAII deposits were distributed around portal regions.

D, E: Immunofluorescence analysis revealed that F4/80-positive Kupffer cells (red) were distributed in close proximity to AApoAII deposits (green).

Figure EV2. TEM images of cells treated with AApoAII at 40 µg/mL for 24 h. Higher magnification images of the interest region (white square) highlights the adhesion of amorphous fibril structures on cell surface.

Figure EV3. Macrophages endocytosed extracellular AApoAII fibrils by macropinocytosis process. Representative confocal images of cells treated with AApoAII at 40 µg/mL and high molecular weight dextran (green) for 24 h, followed by immunostaining using ApoA-II antibody (red).

Figure EV4. The progression of amyloid deposits in pre-CLO mice is independent of AA amyloid and the production of ApoA-II. (A) Representative images of continuous liver sections of Pre-CLO mice stained by ThT (green) and ApoA-II or SAA (red). The amyloid deposits does not contain SAA. (B) Mice were treated with CLO at 25 mg/kg and sacrificed after 1 day, 7 day and 2 months. Serum levels of ApoA-II were determined by western blot analysis (n = 3). (C) Representative immunofluorescence images of amyloid-laden tissues of pre-control and pre-CLO mice using the ApoA-II antibody (green).

